# BioinAI: a general bioinformatic framework for multi-level transcriptomic data analysis using multiple semi-agents

**DOI:** 10.1101/2025.07.21.665890

**Authors:** Mintian Cui, Shixi Wang, Fan Yang, Yifei Wang, Fanyu Kong, Ni Kong, Mengying Li, Xiaoyue Qiao, Zhen Xu, Ziyu Yan, Yu Yan, Jiamo Zhang, Kun Chen

**Affiliations:** Translational Medical Center for Stem Cell Therapy, Institute for Regenerative Medicine, Shanghai East Hospital, School of Life Sciences and Technology, Tongji University, Shanghai 200127, China; Department of Internal Emergency Medicine and Critical Care, Shanghai East Hospital, Tongji University School of Medicine, Shanghai 200120, China; Shanghai Key Laboratory of Signaling and Disease Research, Frontier Science Center for Stem Cell Research, School of Life Sciences and Technology, Tongji University, Shanghai 200092, China

## Abstract

Clinical and biological insights from large-scale transcriptomic data are often limited by technical variability and analytical complexity. Here, we introduce BioinAI, a general bioinformatic framework designed for multi-level transcriptomic data analysis. As part of the BioinAI framework, two algorithms named DeepAdvancer and stNiche were developed to improve data integration and analytical efficiency. DeepAdvancer leverages a class-aware adversarial autoencoder to reconstruct gene-expression profiles from 49,738 samples across 131 skin conditions. These profiles revealed a conserved inflammatory axis and a transcriptomic continuum which link diverse diseases through shared immune responses and structural programs. stNiche leverages spatial graph networks and symmetry-aware matching to identify functional cellular niches and reveals microstructural alterations across development, homeostasis, and disease. For instance, it identified a fibroblast–immune niche surrounding hair follicles in healthy skin, which disappears in pathological states. In addition, BioinAI provides a user-friendly online analysis platform powered by multiple semi-agents (www.bioinai.com), significantly facilitating the extraction of biological insights to advance scientific research.

---

Gene-expression regulation is a complex network, involving many molecular mechanisms and biological functions. Large-scale transcriptome profiling has become a key resource for advancing our understanding of human diseases. Currently, an increasing number of studies aim to gain comprehensive biological insights through integrative analyses of various datasets^1,2^. Conventional integrative methods, such as combat^3^ and limma^4^, are commonly used strategies for removing technical variation. However, these methods, which are typically applied for datasets from the same platform, display a significant decline in performance when applied to cross-platform data analysis^5^. In addition, integrative analysis of large-scale transcriptomic data is often labor-intensive, time-consuming, and requires substantial expertise. Consequently, vast amounts of transcriptomic data remain underutilized. This highlights an urgent need for new approaches to process and integrate large-scale, heterogeneous transcriptomic resources.

In addition to variations in gene expression level, spatial context is also an essential dimension for understanding organ development and disease progression^6^. Therefore, integration of spatial information allows to reveal cellular spatial distributions, offering deeper insights into the tissue microenvironments. Under physiological or pathological conditions, the spatial distribution of cells in the tissue microenvironment often undergoes changes as functional units. Importantly, they should play a vital role in biological processes, such as immune surveillance and response^7^. Currently, approaches to identify the microenvironmental niches mainly rely on similarity clustering, based on the assumption that a niche is a region with homogeneous gene expression within individual samples^8,9^. However, this assumption limits their ability to detect the niches comprising heterogeneous cell populations across samples.

Recent breakthroughs in artificial intelligence (AI) and machine learning are transforming the landscape of bioinformatics^10^, offering new approaches to analyze and interpret complex biological data. Large language models (LLMs)-based intelligent agent systems can autonomously fulfill tasks without constant human intervention by leveraging sequential reasoning, planning, and memory retention^11^. Specifically, the intelligent agent systems are composed of several key modules, including a perception module to receive input, a memory module to retain historical information, a planning module to develop strategies, an execution module to carry out plans, and a communication module to facilitate inter-agent dialogue. Recently, such systems have been applied to bioinformatic analysis. For example, CellAgent^12^ is a type of intelligent agent system designed for autonomously processing and analyzing single-cell RNA sequencing (scRNA-seq) datasets. However, current agent systems continue to suffer from stability and accuracy issues in data analysis, limiting their applicability in rigorous scientific research^11^.

To address these challenges, we developed BioinAI, a comprehensive bioinformatic framework comprising an online platform and two new algorithms, DeepAdvancer and stNiche. DeepAdvancer employs a class-aware adversarial autoencoder to enable efficient integration of cross-platform bulk transcriptomic data, minimizing technical noise while preserving critical biological information. stNiche, leverages spatial graph networks and symmetry-aware matching to identify spatial niches composed of diverse cell types, and further elucidates their functional roles and intercellular communication patterns. In addition, BioinAI integrates advanced LLMs to support omics data analysis, significantly enhancing both the depth and efficiency of transcriptomic research. To ensure robust performance and analytical accuracy, we designed the semi-agent architecture, in which agents are equipped with predefined pipelines, essential code bases, and clearly defined operational roles. This structure strengthens their task-specific reliability and interpretability. Overall, BioinAI provides a precise, efficient, and automated solution for transcriptomic analysis and offers powerful tools for advancing the understanding of disease mechanisms.

## Results

### Design concept of BioinAI

To better interpret gene expression information, we developed the DeepAdvancer, which is designed to reconstruct the biologically meaningful gene expression profiles through weighted combinations of expression profiles from other classes. Within this model, decoder weights are composed to form a matrix whose dimensions correspond to the number of foundational classes multiplied by the number of genes (Methods). This matrix serves as a central expression value for the foundational classes. A loss function is specifically added to minimize discrepancies between this generated matrix and the actual class center values. Class-specific proportional parameters are introduced to finely regulate the contributions of each class. In addition, constraints are applied to the reconstruction loss for the source data, alongside the dataset and class discriminator losses (Fig. 1a). These designs jointly ensure the preservation of intrinsic biological signals while effectively mitigating batch effects.

**Fig. 1:**
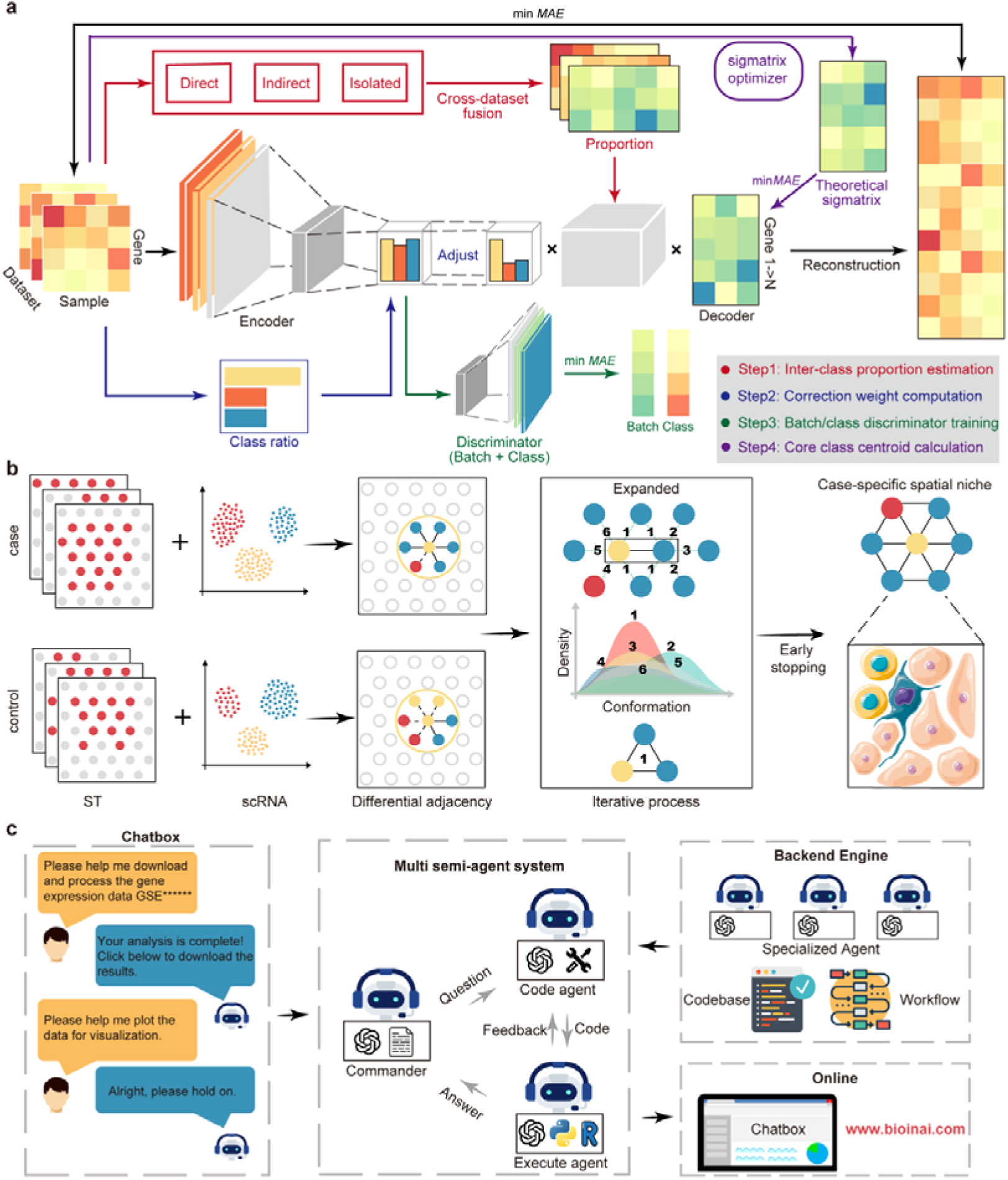
Schematic of the BioinAI Framework. **a**, The DeepAdvancer module integrates and reconstructs bulk transcriptomics datasets across different platforms. It first classifies disease relationships as direct, indirect, or isolated, guiding inter-class proportion calculations. An encoder then extracts features, dynamically adjusted by calculated class proportions. Competitive learning discriminators for class identity and batch identity are applied to remove batch effects and retain biological meaning. Additionally, expression values for foundational classes are optimized from the original datasets through iterative refinement. At last, a distinct optimization ensures minimal discrepancy between reconstructed and source data, thus preserving original biological signatures. **b**, The stNich module identifies biologically relevant cell niches composed of mixed cell types using spatial transcriptomics data. Niches are progressively learned by identifying the conformations of disease-specific cell spots. **c**, A multiple semi-agents conversational system supports user-friendly and stable analysis of transcriptomics data via natural language interaction. The platform is accessible online at www.bioinai.com.

To further investigate spatial architecture, we designed stNiche to identify biologically meaningful functional niches composed of heterogeneous cell types within spatial transcriptomics data. stNiche systematically constructs candidate niches by evaluating structural variations across different biological conditions, such as tissues from diseases and healthy donors. By identifying spatial configurations significantly distinct within these states, the method can isolate niche-level patterns indicative of altered tissue organization. To address the presence of chiral or mirrored spatial structures, the framework incorporates iterative training and symmetry-aware matching. Furthermore, combining cell-type annotations enhances the biological interpretability of detected niches, enabling insights into cell–cell interactions and tissue microenvironments that are context-specific and spatially coherent (Fig. 1b).

In addition, we developed an online platform that enables intuitive and code-free transcriptomics analysis through interactive conversational interfaces. To mitigate hallucinations in large language models, we introduce a semi-agent system. This framework combines automated AI-powered assistance with expert analytical scripts and standardized procedural control, ensuring reliable and reproducible results suitable for rigorous scientific research. Specifically, the platform offers fully automated omics workflows, covering data acquisition, preprocessing, differential expression analysis, enrichment interpretation, and so on (Fig. 1c). In summary, BioinAI offers a comprehensive bioinformatic framework for transcriptomic research.

### DeepAdvancer reconstructs gene expression across 131 skin diseases

Next, we evaluated the performance of DeepAdvancer in integrating large-scale transcriptomic datasets with substantial biological and technical heterogeneity. To this end, we retrieved a collection of gene expression profile datasets related to cutaneous pathologies. Leveraging the automated pipeline developed within the BioinAI platform, we systematically retrieved, annotated, and preprocessed a total of 49,738 samples across 1,016 datasets. These datasets subsequently underwent rigorous filtering and annotation to facilitate downstream visualization and interpretation. The annotated categories included: healthy, autoimmune, cancerous, infectious, allergic, inflammatory, damaged, genetic, aging-related and other (Methods). The majority of the samples were from autoimmune diseases (40.8%), followed by healthy individuals (22.3%) and allergic disorders (13.1%) (Fig. 2a).

**Figure 2:**
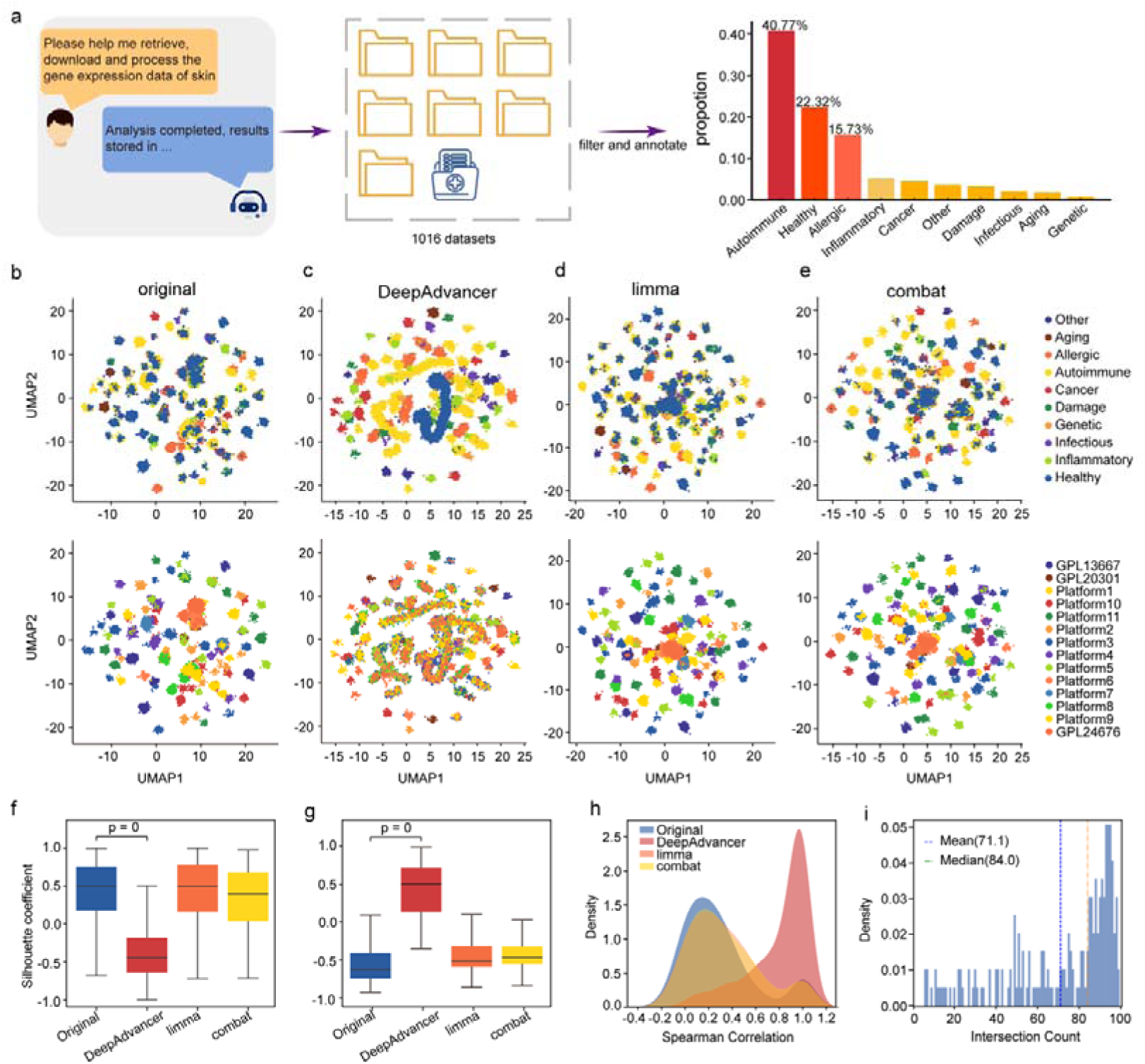
Comparison of batch correction methods on large-scale skin datasets. **a**, Data acquisition and processing were carried out through our BioinAI online platform. The right panel shows the proportion of annotated disease categories. **b–e**, Uniform Manifold Approximation and Projection (UMAP) visualization of the datasets before (**b**) and after batch correction using DeepAdvancer (**c**), Limma (**d**), and ComBat (**e**). The upper row is colored by disease category, and the lower row is colored by dataset platform. **f-g**, Silhouette coefficients by disease class (**f**) and dataset origin (**g**) for each method. **h**, Distribution of spearman correlation for each method. **i**, Histogram showing the intersection count of the top 100 genes ranked by expression fold-change between DeepAdvancer-corrected and reference data.

To assess the efficacy and accuracy of DeepAdvancer, we conducted a rigorous benchmarking exercise. This module was systematically compared against two widely used batch correction methods. The results showed that sample clustering in the original data was primarily driven by batch effects (Fig. 2b). Compared to Limma and ComBat, DeepAdvancer markedly improved the clustering of samples by biological category (Fig. 2c–e). These observations were further supported by quantitative assessments of integration quality (Fig. 2f, g).

To further assess the biological validity of the corrected expression data, we evaluated the accuracy of inter-class expression ratios after batch correction. Consistent with our previous results, the ratios derived from DeepAdvancer-corrected data exhibited the highest concordance with reference ratios (Fig. 2h; Supplementary Fig. S1a). In addition, when comparing the top 100 differentially expressed genes between each pair of classes, DeepAdvancer achieved the highest average overlap (mean = 71.1) with reference gene sets, outperforming limma (mean = 29.7), ComBat (mean = 26.0), and uncorrected data (mean = 24.5) (Fig. 2i; Supplementary Fig. S1b). Furthermore, we conducted a more comprehensive comparison using a smaller psoriasis-focused dataset. Compared to multiple correction methods, including limma, ComBat, Rank-in^13^, Ruaseq^14^, DeepComBat^15^, and MNN^16^, DeepAdvancer consistently demonstrated superior performance (Supplementary Fig. S1c-d).

### Innate immune programs are broadly activated across skin diseases

Using the unified expression data generated by DeepAdvancer, we first identified genes that consistently ranked highly across all skin diseases. These genes were primarily involved in skin inflammation (Fig. 3a, b). Network analysis revealed a core module dominated by innate immune signaling, enriched for hub genes involved in cytokine–receptor interactions (Fig. 3c). To further explore shared pathway-level information, we extracted the pathways captured in the largest number of diseases. As expected, granulocyte chemotaxis, response to bacterial molecules, and inflammatory response were almost universally important (Fig. 3d). This widespread activation shows that skin, as a frontline barrier organ^17^, relies on rapid and nonspecific immune response, underlining a common immune activation program across skin diseases.

**Figure 3:**
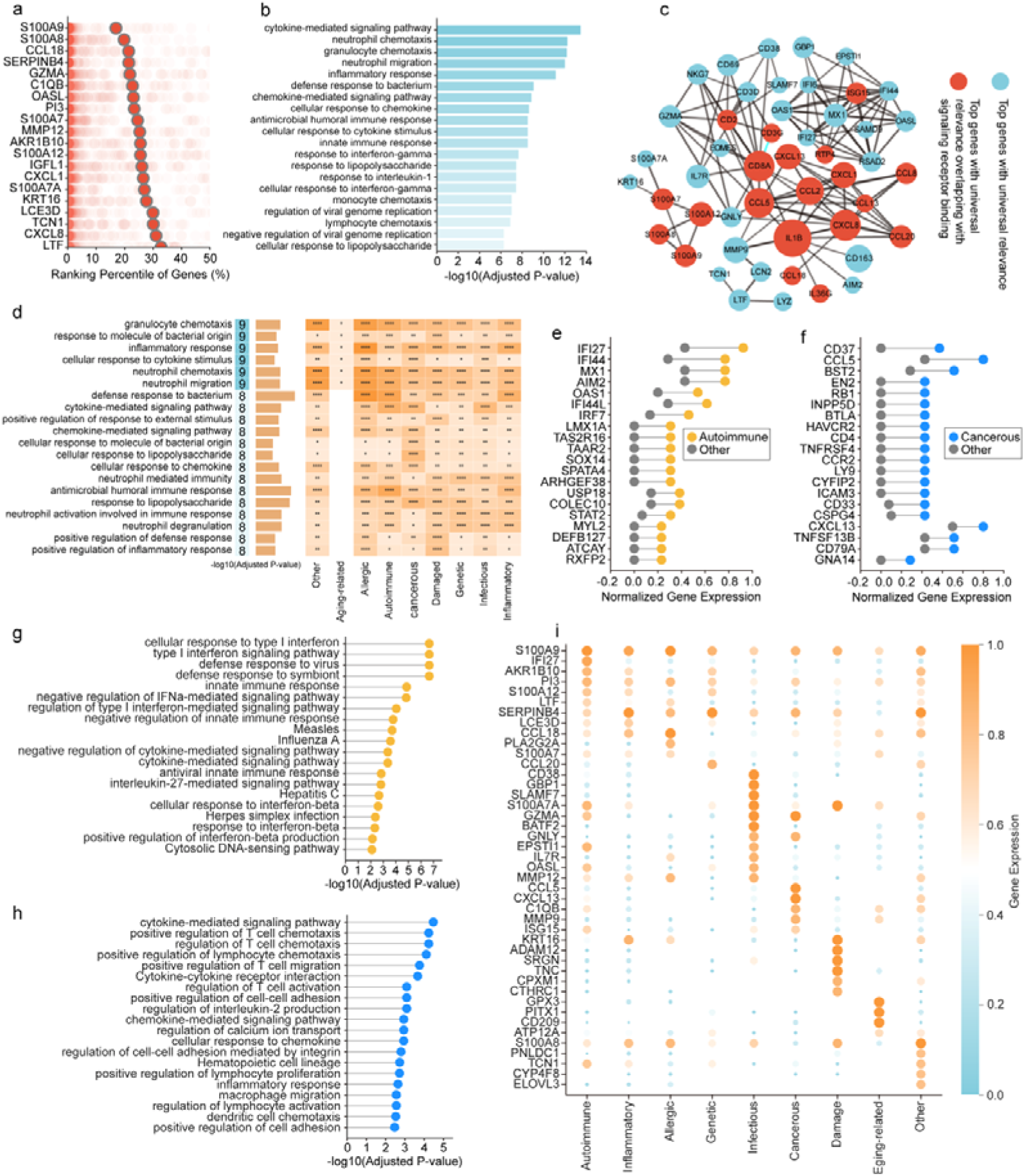
Universally and class-specific important genes and pathways across skin diseases. **a**, List of the highest-scoring genes across all skin diseases. Percentile scores are shown for each class, with average scores highlighted. **b**, The top enriched pathways for the top 100 universally important genes. **c**, Network of top 100 universal important genes. The network was generated with StringDB and disconnected genes are excluded. The size of node was determined by the number of edges. Genes that are included in signaling receptor-related pathways are colored red. **d**, List of top pathways that are universally important. **e**-**f**, The plots of class-specific genes shown for autoimmune class (e) and cancerous class (f). The colored dots show the percentile score of one gene for the specific disease type and the grey dots show the highest percentile score the same gene has among all the other types. The genes are sorted on the basis of the difference values. **g**-**h**, The top enriched pathways for the class-specific genes for autoimmune class (g) and cancerous class (h). **i**, The marker genes for each class.

Next, we aimed to identify genes that only capture variance in specific types. We found that the top autoimmune-specific transcripts were mainly interferon-stimulated genes^18^ (Fig. 3e, g), suggesting that interferon-mediated chronic immune activation may represent a central mechanism underlying autoimmune diseases^19^. By contrast, in skin malignancies, selectively elevated expression of genes was involved in T cell activation, co-stimulatory signaling, and antigen presentation (Fig. 3f–h), underscoring the central role of adaptive immunity within the tumor microenvironment^20^. In infectious skin diseases, DeepAdvancer identified elevated expression of *IRF8*, *GBP1*, *GBP4*, *CD53*, and *IL18RAP* (Supplementary Fig. 2a), indicating that innate immune and antimicrobial responses were activated in these diseases^21^. Top allergic-specific genes (Supplementary Fig. 2b) comprised transcripts associated with chemokines, alongside barrier repair genes, depicting a type 2 immunity–dominated hypersensitivity response^22^. Moreover, genes (Supplementary Fig. 2c) upregulated in inflammatory diseases are involved in MAPK signaling and neuroimmune regulation, suggesting a chronic low-grade inflammatory state. In damaged conditions, strong induction of extracellular matrix (ECM) remodeling genes (Supplementary Fig. 2d) indicated active tissue repair and fibrotic processes^23^. Genetic skin diseases displayed a more complex transcriptional landscape, with class-specific genes (Supplementary Fig. 2e) linked to epidermal structure maintenance, neural signaling, and developmental regulation, suggesting a dysregulation of “structural–neuronal–developmental” axis^24^. In aging-related skin conditions, upregulated genes (Supplementary Fig. 2f) were enriched in oxidative stress responses, mitochondrial metabolism, and cell cycle regulation, reflecting the impact of ROS accumulation and impaired antioxidant defenses^25^. These findings were further supported by marker gene analysis across different types (Fig. 3i, Supplementary Fig. 2g).

### A Transcriptomic Continuum of Skin Diseases

To systematically investigate the relationships among various skin diseases, we performed cluster analysis across all categories (Fig. 4a). We observed that subtypes of the same disease often clustered together, suggesting that DeepAdvancer can capture disease-specific gene expression patterns. Notably, although skin diseases were classified as autoimmune, cancerous, or infectious, the clustering patterns did not strictly follow these predefined categories. A typical example was that psoriasis was closer to actinic keratosis, atopic dermatitis, and squamous cell carcinoma (SCC) than to other autoimmune diseases^26^. We found transcriptional changes across skin diseases occurred on a continuum. This continuum could be broadly divided into three states: (I) a healthy or low-inflammation group characterized by immune regulation and tissue homeostasis (Supplementary Fig. 3a); (II) a high-inflammation group characterized by widespread immune activation (Supplementary Fig. 3b–c); and (III) a tissue destruction group characterized by co-activation of innate and adaptive immune responses, tissue remodeling, and structural breakdown (Supplementary Fig. 3d).

**Figure 4:**
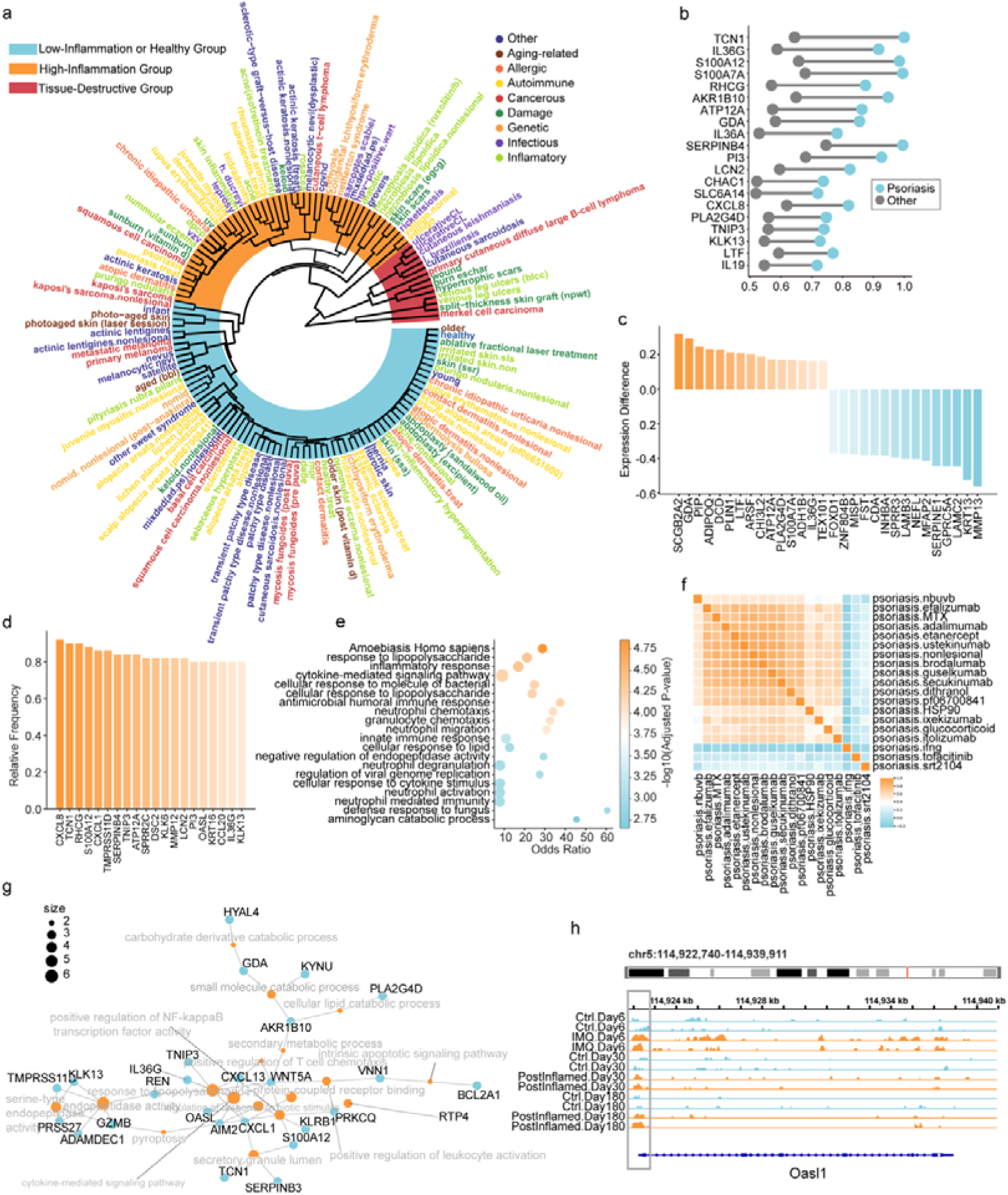
Integrated molecular profiling of skin diseases. **a**, Clustering analysis of various skin diseases based on transcriptomic signatures. **b**, Abnormally upregulated genes associated with psoriasis. **c**, Differentially expressed genes between psoriasis and cutaneous squamous cell carcinoma (SCC). **d**, Genes consistently downregulated across different psoriasis treatments. **e**, Functional enrichment analysis of treatment-associated genes. **f**, Similarity matrix between different therapeutic modalities based on gene expression profiles. **g**, Shared differentially expressed genes between various treatments and the non-lesional skin baseline. **h**, Chromatin accessibility landscape around the *Oasl1* gene locus across different time points.

Interestingly, psoriasis and SCC exhibited similar patterns of transcriptional dysregulation^27^ (Fig. 4b; Supplementary Fig. 3e), yet the disease outcomes were different. We found that, unlike SCC, psoriasis specifically upregulated a set of genes involved in maintaining skin barrier integrity and lipid metabolism^28^. In contrast, SCC upregulated genes were related to extracellular matrix degradation, tumor invasion, and proliferative signaling (Fig. 4c) ^29^. These differences suggest a potential divergence in how keratinocytes respond to and regulate inflammatory stress in the local microenvironment. In psoriasis, keratinocytes remain capable of maintaining barrier function and metabolic homeostasis despite persistent inflammatory stimuli^30^. By contrast, in SCC, inflammatory signaling may drive keratinocytes toward structural instability and invasive reprogramming^31^. Thus, although both conditions are driven by inflammatory pathways, psoriasis represents a non-malignant, adaptive disease state characterized by the coordination of inflammation and barrier reinforcement, whereas SCC follows a trajectory of inflammation-induced remodeling, leading to barrier dysfunction and malignant transformation.

In addition, psoriasis is a prototypical relapsing disease^32^. Patients often experience recurrence after cessation of treatment. To investigate the molecular basis of psoriasis relapse, we analyzed gene expression changes in lesional skin with different therapeutic interventions. We found that although treatment reduced the expression of many inflammation-related genes (Fig. 4d-f), some genes remained elevated compared to unaffected areas^33^ (Fig. 4g). These genes are closely associated with inflammatory response, lipid metabolism, and T cell recruitment. Among them, AIM2 has previously been implicated in the establishment of immune memory following skin damage^34^ (Supplementary Fig. 3f). Intriguingly, we also observed that chromatin accessibility at the *Oasl1* gene^35^ locus remained elevated at day 180 after inflammation resolution (Fig. 4h), suggesting a sustained “inflammatory memory” at the epigenetic level. Targeting these persistent epigenetic programs may be essential for breaking the cycle of psoriasis relapse.

### stNiche reveals niche dynamics across developmental, homeostatic, and diseased human skin

To further investigate how transcriptional programs are spatially organized in the skin, we analyzed ST profiles from 240 samples (Fig. 5a). Uniform Manifold Approximation and Projection (UMAP) projection revealed 17 distinct cellular subpopulations (Fig. 5b) across various pathological states (Fig. 5c, d). To further achieve spatially resolved cell-type localization, we applied Cell2location^36^ to decompose cell-type composition across ST spots via annotated scRNA-seq references. This enabled us to reconstruct the spatial distribution of distinct cell populations within each tissue section (Fig. 5e-g; Supplementary Fig. 4a).

**Figure 5:**
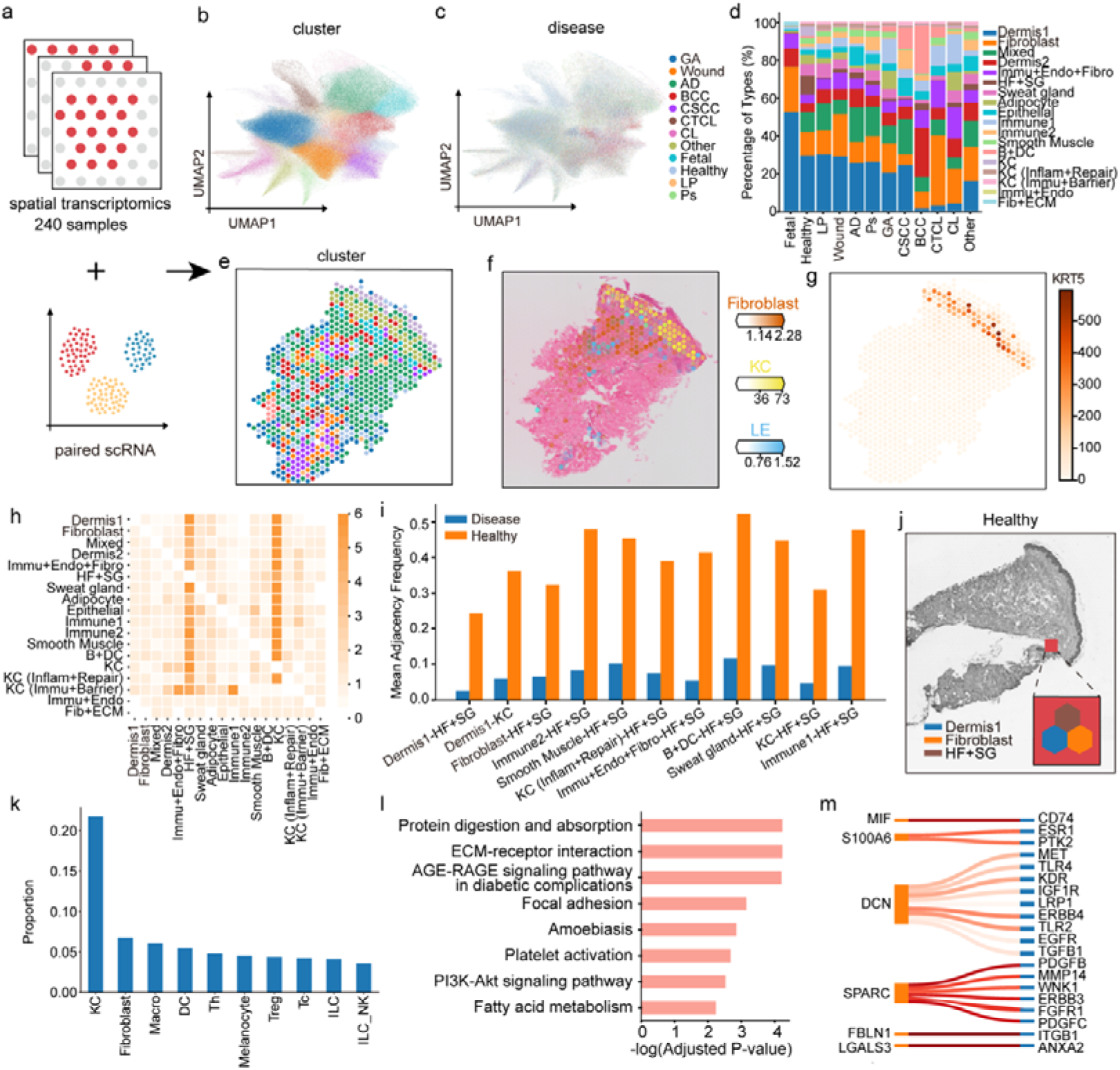
Identification and characterization of spatial niches in human skin using stNiche. **a**, Schematic overview of the integration of ST and matched single-cell RNA-seq datasets. b, UMAP visualization of distinct cellular clusters. **c**, Distribution of cellular subpopulations across various skin conditions. **d**, Stacked bar plot showing changes in cellular composition across different diseases. **e–g**, Cell-type deconvolution of ST spots (e) using Cell2Location based on scRNA-seq annotations, with spatial mapping of inferred cell types (f) and corresponding marker gene expression (g). **h**, Heatmap showing differences in second-order structures between healthy and diseased samples. **i**, Bar plot showing significantly altered second-order spatial interactions. **j**, Spatial niche identified by stNiche, composed of three co-localized spot types enriched for hair follicle, fibroblasts, and immune cells. **k**, Proportions of major cell types within the identified niche. **l**, Enriched functional pathways associated with the niche. **m**, Ligand–receptor interaction network highlighting potential signaling between the niche and surrounding microenvironment.

Next, we investigated disease-associated alterations in spatial tissue organization using stNiche (Methods), and found that several second-order structural connections were significantly reduced in disease samples compared to healthy controls (Fig. 5h–i). Notably, stNiche identified a niche comprising three spatially co-localized spot types enriched for hair follicle, fibroblasts, and immune cells (Fig. 5j, k). Functional enrichment analysis revealed significant enrichment in ECM–receptor interaction^37^, AGE–RAGE signaling^38^, focal adhesion, and the PI3K–Akt pathway^39^ (Fig. 5l). Furthermore, ligand–receptor interaction analysis of stNiche uncovered a signaling network centered around *MIF*, *S100A6*, *DCN*, and *SPARC*, which interact with cell-surface receptors such as *CD74*, *ESR1*, and *TGFBR1* (Fig. 5m). These interactions are implicated in driving epithelial repair, matrix reorganization, and local inflammatory activation.

We further applied stNiche to psoriasis samples and identified a niche composed of dermis1 and immune2 cell clusters (Supplementary Fig. 4b). Functional enrichment analysis revealed that this niche was enriched in lipid metabolism^40^ and inflammatory pathways (Supplementary Fig. 4c), consistent with our prior findings (Fig. 4g). In basal cell carcinoma (BCC), we identified a niche consisting of dermis2 and ‘Immu+Endo+Fibro’ cell clusters (Supplementary Fig. 4d), enriched for cancer-associated and endocrine/metabolic pathways, suggesting a multifunctional role in immune evasion (Supplementary Fig. 4e). In fetal skin, a niche formed by dermis1 and fibroblasts (Supplementary Fig. 4f) was enriched for pathways related to ECM remodeling, PI3K–Akt signaling, and cell adhesion and migration^41^, pointing to its involvement in structural development and local tissue organization. Together, these findings demonstrate that spatial niches exhibit high state-specificity across developmental, homeostatic, and pathological conditions, and that niche remodeling may serve as a key driver of disease progression.

### BioinAI provides a comprehensive analysis platform for Bulk RNA-seq data

In recent years, transcriptomics has emerged as a crucial tool for elucidating disease mechanisms and identifying therapeutic targets^42^. However, its analytical process remains complex and time-consuming. To address this, we developed the BioinAI online module (hereafter referred to as BioinAI-Web) that enables automated transcriptomic analysis through a conversational interface (Fig. 6a). Users can initiate analysis workflows, upload datasets, and download results through natural language commands (Supplementary Fig. 5a). Specifically, BioinAI-Web supports automated dataset retrieval and preprocessing. Based on the experimental design and data type, the system intelligently determines appropriate normalization procedures and gene symbol conversions (Supplementary Fig. 5b), generating standardized gene expression matrices and corresponding phenotype tables (Fig. 6b). During analysis, BioinAI-Web accommodates a range of common transcriptomic tasks (Fig. 6c). For instance, the platform automatically performs differential expression analysis, with volcano plots illustrating the distribution of differentially expressed genes (Fig. 6d–e). In addition, boxplots allow intuitive comparisons of key gene expression between groups (Fig. 6f). The platform also integrates a suite of downstream analytical modules, including functional enrichment analysis (Fig. 6g), gene set enrichment analysis (Fig. 6h), weighted gene co-expression network analysis (Fig. 6i, Supplementary Fig. 5c, Fig. 6j), and PPI network reconstruction (Fig. 6k).

**Figure 6:**
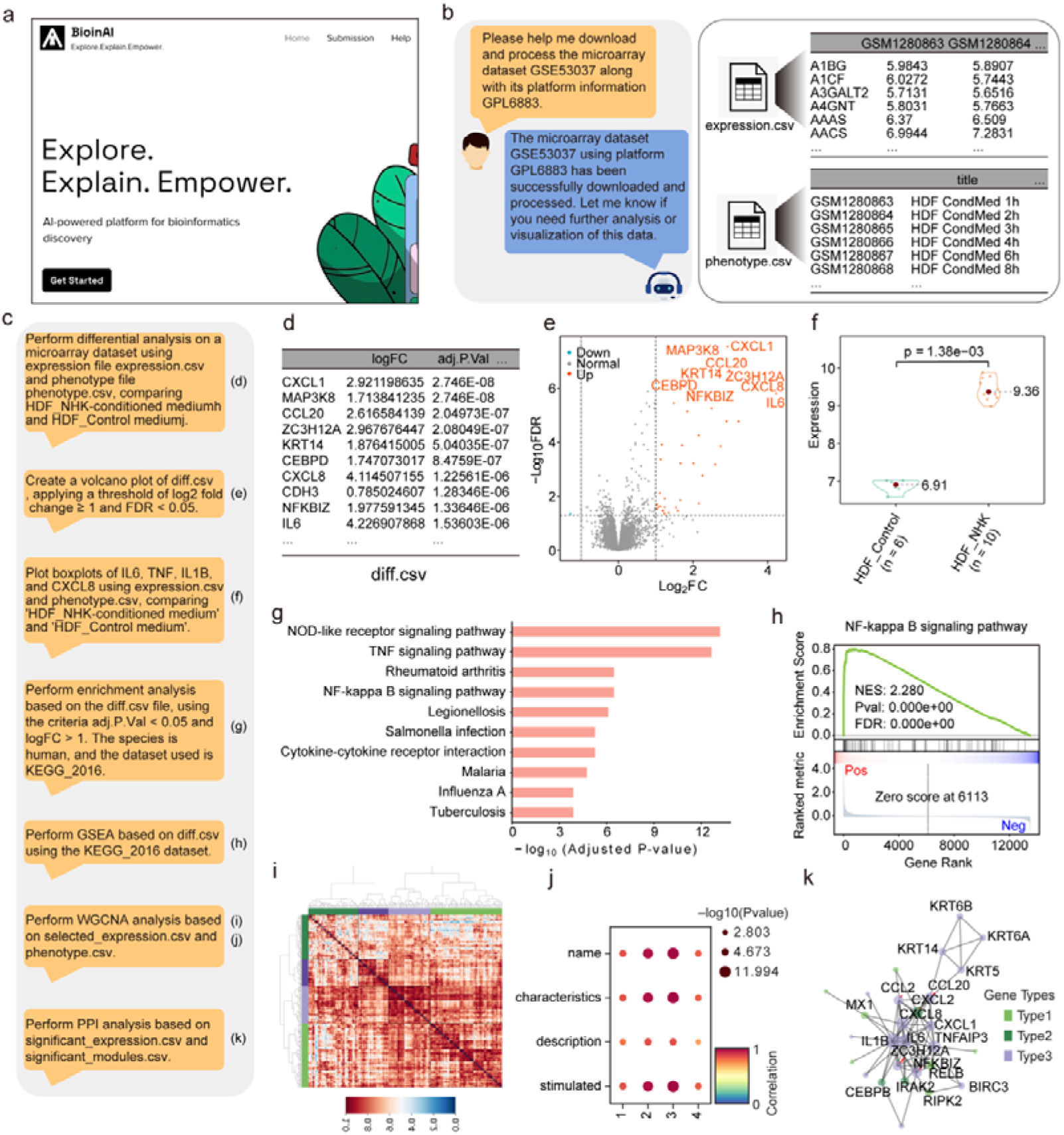
BioinAI-Web enables automated analysis of bulk transcriptomic data. **a**, Screenshot of the BioinAI online platform. **b**, Example conversation illustrating a user request to download and preprocess transcriptomic data (left), with the resulting standardized gene expression matrix and phenotype metadata (right). **c**, Sequence of user prompts and corresponding results. **d**, Output of differentially expressed genes. **e,** Volcano plot showing the distribution of significantly upregulated and downregulated genes. **f**, Box plot comparing expression levels of representative inflammation-related genes across two groups. **g**, Bar plot of enriched functional terms among upregulated genes. **h**, Gene set enrichment analysis highlights significant upregulation of the NF-κB pathway. **i**, Heatmap displaying weighted gene co-expression network analysis (WGCNA) module assignment for differentially expressed genes. **j**, Correlation between gene modules and clinical traits. **k**, Protein–protein interaction network of top differentially expressed genes.

To evaluate the utility and performance of BioinAI-Web, we conducted two exemplary analyses using the microarray dataset GSE53037^43^ and the RNA-seq dataset GSE114729^44^. In GSE53037, differential expression analysis revealed significant upregulation of genes associated with inflammation and cell migration in fibroblasts co-cultured with keratinocytes^45^ (Fig. 6e–f). Functional enrichment analysis of these genes highlighted canonical pathways such as NOD-like receptor, TNF, and NF-κB signaling. (Fig. 6g–h). Co-expression network analysis identified four gene modules (Fig. 6i), among which module 3 showed the strongest association with the co-culture condition (Fig. 6j) and included key genes such as *COPS3*, *SIRPA*, and *TNFAIP2* (Supplementary Fig. 5c). PPI analysis further revealed that these inflammatory mediators formed a densely interconnected signaling network (Fig. 6k). In GSE114729, BioinAI-Web generated standardized expression matrices and phenotype annotations across pre-treatment, post-treatment, and relapse stages of psoriasis. Comparative analysis revealed significant upregulation of lipid metabolism genes in relapse^46^ (Supplementary Fig. 5d–e; Fig. 4g). Pathway enrichment showed treatment suppressed pro-inflammatory pathways including NOD-like receptor and cytokine receptor interactions, while enhancing drug metabolism pathways (Supplementary Fig. 5f–i). However, these inflammatory pathways became reactivated after drug withdrawal (Supplementary Fig. 5h, 5j).

### BioinAI provides a comprehensive analysis platform for scRNA-seq and ST data

BioinAI-Web also supports intelligent analysis of single-cell and spatial transcriptomics datasets (Fig. 7, Supplementary Fig. S6). Users can initiate the analyses simply by submitting natural language commands (e.g., “Analyze scRNA dataset GSE220116^47^” or “Process spatial transcriptomics dataset Tildra^7^”). The system will then automatically launches the corresponding workflows. For single-cell datasets, BioinAI-Web executes a modular pipeline that includes quality control, batch correction, dimensionality reduction, clustering, cell type annotation, subpopulation proportion analysis, differential expression analysis, and cell–cell communication inference. If users wish to define custom experimental groups, they can upload grouping files along with the data, and the platform will automatically adapt to perform stratified analyses.

**Figure 7:**
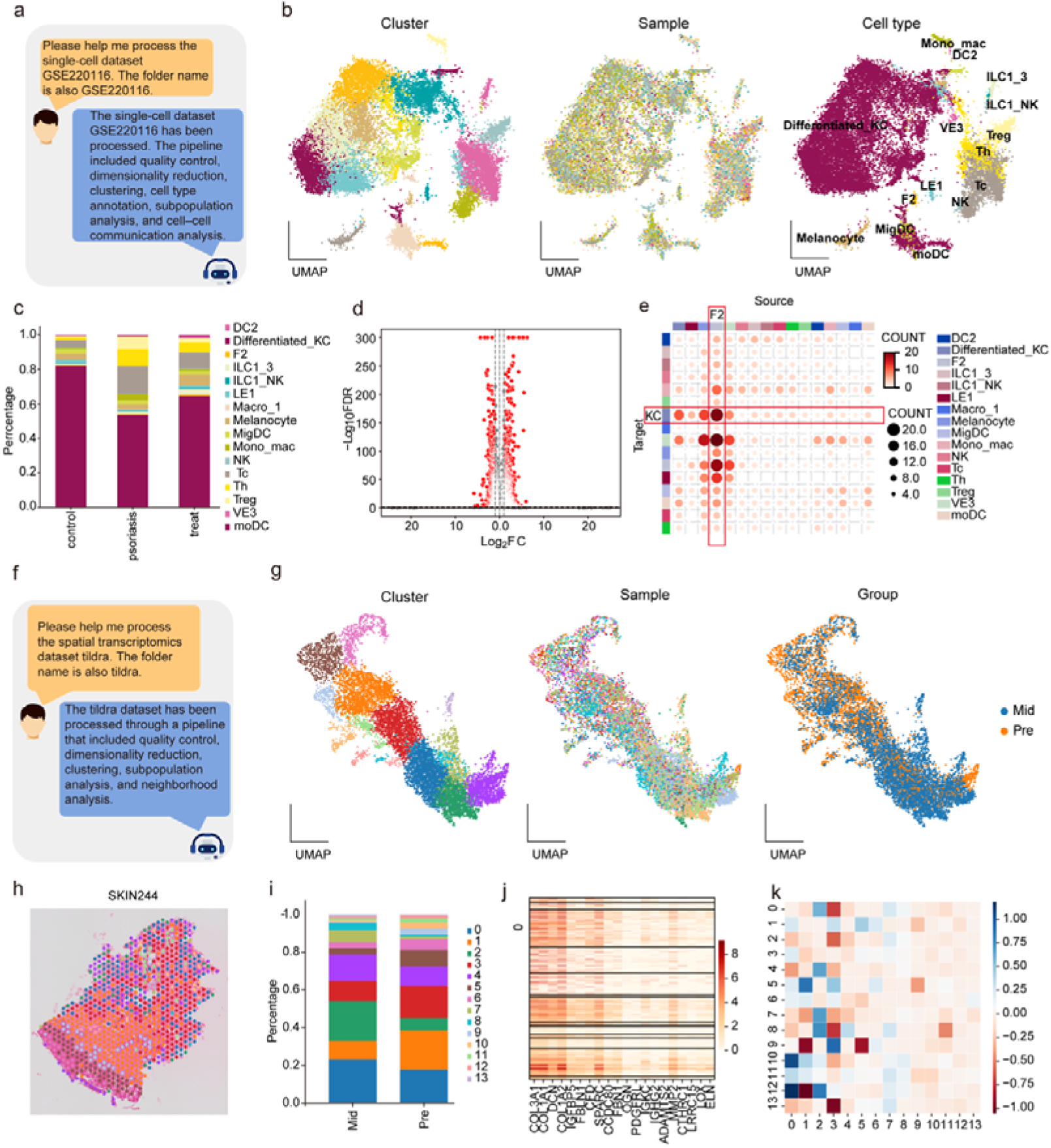
BioinAI-Web enables analysis of single-cell and spatial transcriptomics data. **a**, Representative query to the online platform requesting single-cell RNA-seq data processing and the corresponding response. **b**, UMAP plots showing integrated single-cell data colored by cluster (left), sample (middle), and annotated cell types (right). **c**, Stacked bar plots illustrating the proportion of cellular subpopulations across different groups. **d**, Volcano plot showing differentially expressed genes in the keratinocyte subpopulation between healthy individuals and psoriasis patients. **e**, Heatmap depicting intercellular communication changes among subpopulations in psoriasis. **f**, Representative query to the online platform requesting spatial transcriptomics data processing and the corresponding response. **g**, UMAP plots of integrated spatial transcriptomics data colored by cluster (left), sample (middle), and group (right). **h,** Spatial mapping of subpopulations in psoriasis skin (sample Skin244). **i**, Stacked bar plot showing changes in subpopulation composition before and after treatment. **j**, Heatmap of marker gene expression in subpopulation 0. **k**, Heatmap illustrating adjacency changes between subpopulations pre- and post-treatment.

As a practical example, we analyzed dataset GSE220116, which includes single-cell profiles from psoriatic skin before and after treatment, along with healthy controls. BioinAI-Web integrated the data and constructed a UMAP embedding colored by cluster, sample origin, and annotated cell types (Fig. 7b). Multiple subpopulations were identified, including keratinocytes, fibroblasts, and immune cells (Fig. 7c). Differential expression analysis was performed across all subtypes by BioinAI-Web (Fig. 7d). Cell–cell communication analysis revealed markedly enhanced interactions between ‘KC’ and ‘F2’ clusters in psoriatic skin^48^ (Fig. 7e, Supplementary Fig. S6c), which were notably reduced after treatment (Supplementary Fig. S6d). These interactions were centered around pro-inflammatory ligand–receptor pairs, including CXCL14_CXCR4 and CD44_TYROBP (Supplementary Fig. S6e).

For spatial transcriptomic analysis, we demonstrated the platform using the Tildra dataset. BioinAI-Web automatically performed quality control, dimensionality reduction, clustering, and neighborhood graph analysis (Fig. 7g). It visualized the spatial distribution of clusters in a psoriasis patient (Fig. 7h) and quantified changes in spatial subpopulation composition before and after treatment (Fig. 7i). Further marker gene and spatial adjacency analyses (Fig. 7j–k) revealed spatial remodeling of fibroblast-associated clusters during treatment. Together, these results demonstrate that BioinAI-Web can efficiently handle not only bulk transcriptomics but also complex single-cell and spatial transcriptomic data.

## Discussion

In this study, we developed BioinAI, an intelligent, unified, and efficient framework for multi-dimensional transcriptomics analysis, comprising an online platform and two new methods, DeepAdvancer and stNiche. Through the analysis of large-scale datasets, we demonstrate the robustness of these two methods in cross-platform data integration, biological signal extraction, and spatial niche deconstruction. In addition, the BioinAI online platform provides a user-friendly and fully-automated analysis ecosystem, streamlining the entire workflow from data retrieval and preprocessing to downstream interpretation.

DeepAdvancer establishes a unified embedding space for bulk RNA-seq data across diverse platforms. Compared with traditional methods, it captures disease-relevant features more effectively under high heterogeneity. Applying DeepAdvancer to large-scale skin datasets, we uncovered a transcriptomic continuum of skin diseases characterized by a gradient of inflammatory activation^49^. Notably, psoriasis and SCC shared similar dysregulated immune signatures, including chemokine activation and epithelial remodeling^26,27^. However, psoriasis retained barrier-related gene expression, suggesting an “adaptive inflammation” state^30^ — a non-destructive adaptation to sustained immune activation pressure. In contrast, SCC showed upregulation of invasion-associated genes, indicating a shift toward a “remodeling-invasion” axis^29^. Overall, our method enables a unified transcriptomic analysis of the entire dataset, offering new insights from a more comprehensive perspective.

stNiche applies a spatial graph learning framework to identify functionally coordinated cellular niches across tissue states. In healthy skin, we identified a follicle-associated niche^50^ composed of immune cells, fibroblasts, and epithelial components. This niche is enriched for signaling pathways related to stromal remodeling, epithelial repair, and immunomodulation. We think it might function as a spatially organized unit for modulating tissue integrity by immune regulation from local microenvironment. In disease contexts such as psoriasis and BCC, we observed distinct architectures with altered functions^51^. These findings suggest that disease progression involves not only cell-type changes but the disintegration and reprogramming of spatially defined microenvironments, highlighting niche remodeling as a key hallmark of pathogenesis.

Our large-scale data mining efforts were enabled by the BioinAI-Web, which integrates a semi-agent architecture with large language models. This design supports natural-language interaction, modular workflow execution, and stable outputs, while mitigating common issues such as LLM hallucinations^11^. Compared with fully model-driven systems^52^, BioinAI-Web offers a complementary approach that balances automation with interpretability and analytical control. Benchmark analyses demonstrate the platform’s scalability and versatility in integrating high-throughput datasets and extracting biological insights. Future work will focus on enhancing multi-omics integration, improving agent–user interactions, and incorporating more advanced analytical algorithms. Together, BioinAI-Web represents a step toward the automation, standardization, and intelligent transformation of transcriptomic research. It provides a powerful tool for dissecting disease mechanisms and paves the way for next-generation scientific exploration.

## Online Methods

### Collection of Bulk Transcriptomic Data via BioinAI-Web

Bulk transcriptomic datasets related to human skin were retrieved and preprocessed using BioinAI-Web, a multi-agent system designed for bioinformatics analysis. The query command “retrieve GEO datasets for human skin transcriptomes” was issued to BioinAI, which automatically identified and downloaded relevant RNA-seq and microarray datasets from GEO. A subsequent command “download and preprocess RNA-seq/microarray expression data” triggered standardized parsing, normalization, and formatting of raw expression matrices. All datasets were stored in a predefined directory and converted into standard gene expression matrices. Quality control was performed in multiple stages. Datasets were excluded if they were derived from cell lines, contained incomplete expression matrices, lacked sample-level annotations, or generated on undefined platforms. For the retained datasets, biological context metadata, such as tissue origin, disease condition, and treatment status, were extracted and harmonized into a unified metadata table for downstream analysis.

To ensure cross-dataset comparability, all expression matrices were harmonized using unified gene symbols. Genes with high-missing rates across datasets were removed. Datasets with abnormally low gene counts, likely due to low probe coverage or platform incompatibility, were excluded to improve data quality. To reduce technical noise and enhance downstream analytical robustness, normalization steps were applied. Extreme expression values were clipped, log-transformation was performed where appropriate, and redundant samples with identical expression profiles were removed. For missing values, within-sample mean imputation was used. The final integrated bulk transcriptomic dataset spans a wide range of skin conditions, including healthy tissue, inflammatory disorders, autoimmune diseases, and skin cancers. This resource served as the foundation for subsequent multi-task analysis and model training.

### Pre-training data preparation

To systematically quantify gene expression differences between reference classes and other disease classes, we developed and applied the BioinAI_FC module—a tool optimized for efficient logFC estimation across heterogeneous bulk transcriptomic datasets. The module employs a hierarchical strategy based on the degree of class co-occurrence within datasets and divides pairwise class relationships into three types: directly shared, indirectly shared, and non-overlapping. For directly shared pairs, in which both classes appear within the same dataset, the corresponding samples are extracted to construct design and contrast matrices. Differential expression analysis is performed using the limma package, including log2 transformation, outlier clipping, model fitting, and empirical Bayes statistics. For indirectly shared pairs, in which the two classes do not co-occur in the same dataset but both share an intermediate third class. The median expression of the intermediate class is used to bridge the two datasets. Batch effects are removed using limma-based correction prior to logFC computation. For non-overlapping pairs, in which no shared class exist, datasets containing the individual classes are merged. After preprocessing and transformation, limma is also used to estimate differential expression after batch correction. This strategy is considered less robust due to potential bias and is used only when no shared classes are available. To improve robustness, comparisons from multiple datasets are filtered based on expression profile similarity, and representative subsets are selected for integration. The final result is a multi-class logFC matrix serving as input for downstream tasks.

Prior to model training, the expression profile of baseline classes was further optimized to improve cross-class feature consistency. Initial baseline values were computed as the mean expression of the base classes across datasets. An optimization algorithm was then applied to refine the baseline profile using information from directly shared classes. Specifically, one-hop shared classes that co-occurred with the base class in at least one dataset were first identified. Corresponding logFCs between the base class and these shared classes were extracted. Then, the initial expression matrix was optimized to maximize expression pattern similarity of the same one-hop class as calculated from different baseline classes. Two types of sharing relationships were considered: (1) one-hop classes shared by all baseline classes, and (2) pairwise sharing between subsets of baseline classes. To prevent overfitting and drift from the original center, an L2 regularization term was added to constrain deviations from the initial mean vector. Optimization was performed via gradient descent with adaptive scheduling of the learning rate and regularization coefficient. This yielded a stable and biologically interpretable center matrix.

To retain full gene expression information and support downstream modeling, the raw gene expression matrix was used directly as model input, without applying dimensionality reduction or feature selection. To improve inter-sample comparability and mitigate biases arising from differences in numerical scale, MinMax normalization was applied to each dataset independently, scaling gene expression values to the range [0,1]. This design encourages the model to learn stable representations and reduces gradient instability.

To improve the balance of training data, all samples were grouped into 14 platform categories based on technological similarity. Platforms with minimal sample counts or structural redundancy were merged to ensure adequate representation across categories. To enhance the interpretability and generalizability of downstream disease-level analysis, disease samples were further annotated into eight major clinical groups based on etiological mechanisms or shared clinical features. These included: (1) autoimmune diseases (e.g., systemic sclerosis, lupus erythematosus); (2) allergy-related disorders (e.g., atopic dermatitis, chronic urticaria); (3) infectious diseases (bacterial, viral, or fungal infections of the skin); (4) genetic disorders (congenital or hereditary skin diseases); (5) injury-related conditions (e.g., trauma, burns, postsurgical healing); (6) aging- and environment-associated conditions (e.g., photodamage, aging); (7) cancers (primary cutaneous carcinomas, skin lymphomas); (8) inflammatory conditions not clearly attributable to immune or infectious origins but showing pronounced skin inflammation; (9) others, including samples with ambiguous or undefined diagnostic categories. This disease reclassification was used for visualization only and was not involved in model training.

### BioinAI_DeepAdvancer framework

The DeepAdvancer framework is a gene expression decomposition model that enables the reconstruction of expression profiles for any class using a weighted combination of profiles from other classes. Let *X*∈*R^N^*^×*M*^ represent the input gene expression matrix, where *N* is the number of samples and *M* is the number of genes. To capture class-specific contributions, we introduce a class-specific weighting tensor *P*∈*R^P^*^×*C*×*M*^, where *F* denotes the latent feature dimension, *C* is the number of classes, and the weights modulate the contribution of each class to the overall gene expression profile. The model is implemented using an autoencoder architecture. The encoder network *f_ϕ_*:*X*→*Z* maps the input data *X* to a latent representation Z∈R*^D^*, while the decoder network *g_Ψ_*:*Z*→*X* maps the latent representation back to the original expression space. The training objective is to minimize the reconstruction loss between the input and the reconstructed expression: In addition, we define a decoder transformation matrix *W*∈R*^D^*^×*M*^, which represents the expression centers of the base classes in the reconstructed space. To ensure that the learned decoder preserves class-specific structure, we introduce a

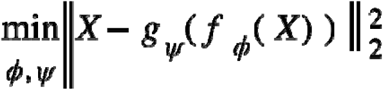

regularization loss that penalizes the deviation between and the true class-wise expression centers :

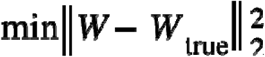

To improve model robustness and mitigate batch effects, we further introduce two adversarial objectives: a batch discriminator loss *L*, and a class discriminator loss *L*, The full training objective is therefore formulated as:

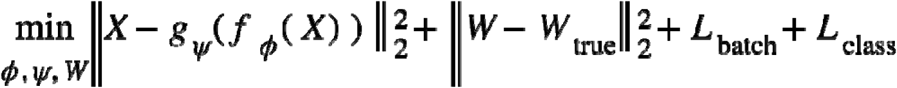

This architecture ensures that the model not only effectively removes batch-related effects but also preserves the original expression characteristics of the data.

### Batch effect correction using multiple methods

To benchmark DeepAdvancer against other correction strategies, we applied a range of established batch effect correction methods to the input expression matrices. These methods span linear models, nonlinear mappings, and deep learning–based frameworks, representing diverse theoretical approaches to batch effect mitigation. The following methods were used: (1) ComBat, an empirical Bayes–based linear model widely applied in both microarray and RNA-seq studies. We incorporated disease categories as covariates in the design matrix to preserve biological variability while removing technical effects. (2) removeBatchEffect from the limma package, a fast linear adjustment method that fits gene-wise linear models and removes batch-associated components. This served as the default correction strategy in our primary modeling pipeline. (3) RUVSeq, which infers and removes unwanted variation using negative control genes or a set of presumed invariant genes. This method is particularly suited to RNA-seq datasets lacking explicit batch annotations. (4) Rank-in, a normalization method based on expression rank preservation across sample groups. It is robust in cross-platform correction tasks. (5) MNN (mutual nearest neighbors), which performs nonlinear alignment of samples by identifying mutual neighbors in latent expression space. (6) DeepComBat, a hybrid method that combines the feature extraction capabilities of deep neural networks with the statistical correction strengths of ComBat. For all methods, we ensured a consistent gene space and harmonized sample annotations. Correction outcomes were systematically evaluated under a unified framework.

### Evaluation of batch correction performance

To systematically assess the effectiveness of different batch correction methods, we performed both structural and quantitative evaluations on the corrected expression matrices using clustering consistency scores. First, we evaluated the compactness and separation of sample groups in the reduced-dimensional space using either disease labels or batch labels. For each class, we computed a class-specific centroid and calculated two types of distance-based metrics for each sample: (1) the average distance to other samples of the same class, and (2) the average distance to the ten nearest centroids of other classes. These values were then used to compute a silhouette score, defined as:

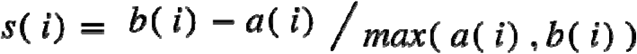

where *a*(*i*) is the average nearest *b*(*i*) inter-class distance. To further assess biological consistency, we computed the similarity between predicted and true logFCs for each one-hop shared class pair after correction. In addition, we evaluated the overlap of the top 100 differentially expressed genes before and after correction to assess the fidelity of signal retention.

### Downstream analysis of corrected expression data

To identify common genes, we first extracted gene expression profile of all healthy samples. For each disease class, differential expression was then calculated relative to this healthy baseline. Within each disease group, we ranked genes by their differential expression and recorded the top genes per disease. High-frequency genes were defined as those recurrently appearing in the top lists across multiple diseases within the same group. To evaluate the consistency of expression trends for these high-frequency genes across all disease classes, we ranked differential expression genes within each class. We then performed functional enrichment analysis on the top high-frequency genes. The top enriched pathways were visualized. To identify disease-specific genes, we computed a specificity score for each gene by taking the difference between its rank in the target class and its highest rank in all other classes. Genes with the largest positive differences were defined as specific to that disease. Functional enrichment was also performed on these disease-specific gene sets, and the top-ranked genes per disease class were visualized separately.

### Collection and processing of scRNA-seq and spatial transcriptomics data

ST datasets of human skin tissue were retrieved through systematic database searches, along with matched scRNA-seq data from corresponding tissue samples. For ST data, batch effect correction was first applied to integrate samples. The data were then normalized, dimensionally reduced, and clustered to identify and define spatially distinct regions based on transcriptional signatures. For the paired scRNA-seq data, standard preprocessing was performed, including normalization, quality control, and dimensionality reduction. Cell type annotation was conducted using well-established marker genes, resulting in a high-confidence cell-type reference dataset. Cell type decomposition of the ST data was performed using Cell2location, a probabilistic model that leverages annotated scRNA-seq reference data to estimate the cell-type composition of each spatial spot. This approach enabled fine-resolution spatial mapping of cell types across the tissue section.

### Spatial niche identification using stNiche

We developed stNiche, a framework for identifying tissue microenvironments based on ST data. First, the adjacency relationships of spot cluster were extracted to construct a sample-specific adjacency matrix. Adjacency was defined between each spot and its immediate spatial neighbors. For any pair of cell cluster types *L_x_* and *L_y_*, we computed the frequency of their adjacency 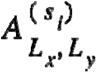 within each sample

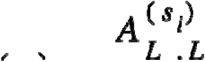

*s_i_*, and normalized it to obtain an adjacency proportion:

where 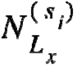 denotes the total number of *L_x_* in *s_i_*. We then assessed group-level differences in adjacency proportions and evaluated statistical significance using the Mann–Whitney U test, followed by Benjamini–Hochberg correction for multiple comparisons, resulting in a final significance matrix.

Based on statistically significant adjacency pairs, we defined higher-order structural units. Starting from pairwise (second-order) neighborhoods, we expanded these

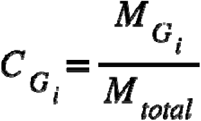

structures by iteratively including neighboring spots of each current position, provided that was not already in the structure and belonged to the same spatial sample. Each identified structure was then assigned a geometric signature, defined by the relative spatial coordinates among its spots, allowing structure-level comparison and classification. Two structures *S_a_* and *S_b_* were assigned to the same group if their relative geometries and cluster labels were fully matched. For each structure group *G_i_*, we computed its coverage ratio:

where 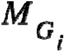 is the number of samples in which structure group *G_i_* is observed, and *M_total_* is the total number of samples. To identify enriched structures, we calculated FC in group-specific frequencies:

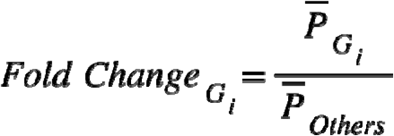

where 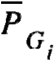 is the average frequency of *G_i_* in the target group and 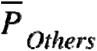 is the average of other structures.

We retained structures with FC above a predefined threshold and minimum coverage, and applied an iterative expansion strategy to refine structure boundaries. In each round, expanded structures were re-evaluated for significance and coverage. If expansion failed to meet criteria, the previous valid structure was retained. This hierarchical and statistically grounded strategy enables robust identification of spatial niches that reflect biologically meaningful microstructures in tissue architecture.

### Functional and signaling analysis of spatial niches

After identifying spatial niche structures, we further characterized their cellular composition and functional relevance. Based on prior high-confidence cell type annotations from scRNA-seq, we determined the cell type composition within each niche region by Cell2location. To investigate niche-specific functional activity, we performed differential gene expression analysis between niche regions and non-niche spots from the same tissue samples. Differentially expressed genes were identified as those significantly expressed in niche regions. Functional enrichment analysis was then conducted on these genes to reveal the biological processes and pathways associated with each niche type. To further explore communication between niches and their surrounding microenvironment, we used the Squidpy^53^ framework to infer spatial signaling interactions. Specifically, we assessed ligand–receptor activity between niche spots and adjacent spatial domains, enabling the reconstruction of spatially resolved intercellular communication networks.

### Construction of the BioinAI online platform

To enable automated processing and intelligent analysis of bulk transcriptomic data, we developed the BioinAI online platform (accessible at www.bioinai.com) based on the AgentScope framework^54^. The platform integrates multiple semi-agent collaboration with LLM interfaces to support natural language–driven omics analysis. At the core of the backend system is a ReActAgent architecture, which orchestrates key analytical tasks including data retrieval, preprocessing, differential expression analysis, feature integration, and so on. The frontend adopts a modular design, offering functionality for data upload, results download, and user support. Users interact with the platform through natural language prompts. The system interprets these requests and automatically executes the corresponding analysis pipeline, including matrix construction, normalization, model inference, and result generation. BioinAI-Web currently supports complete pipelines for bulk transciptomics data, single-cell, and spatial transcriptomic data.

## Data availability

The main data supporting the results of this study are available within the paper and its Supplementary Information. Input gene expression datasets, reconstructed expression profiles, and integrated spatial transcriptomics datasets are available on Figshare at https://doi.org/10.6084/m9.figshare.2958995955.

## Code availability

The code necessary to reproduce our findings is available in GitHub at https://github.com/BioinAI.

## Acknowledgements

The work was financially supported by National Natural Science Foundation of China (Nos. 32370961), Shanghai Rising-Star Program (No. 21QA1407600), Open Research Fund of Basic Medicine College (No. JCKFKT-ZD-004), Fundamental Research Funds for the Central Universities (No. 22120250158, 22120250374).

## Author contributions

K. C. designed and supervised the research. M. C., and S. W. organized and analyzed the data. M.C. developed the framework, created the figures, and drafted the manuscript. F.Y. and Y.W. proofread and revised the manuscript. M.C., F.K., N.K., M.L., X.Q., Z.X., Z.Y., Y.Y., and J.Z. participated in the development and testing of the online platform. All authors read, verified the data, and approved the final version of the manuscript. K. C. was responsible for the decision to submit the manuscript.

