## Supplementary Materials for "BioinAI: a general bioinformatic framework for multi-level transcriptomic data analysis using multiple semi-agents"

**This PDF file includes:**

Supplementary Figures 1 to 7.

**Supplementary Figures**

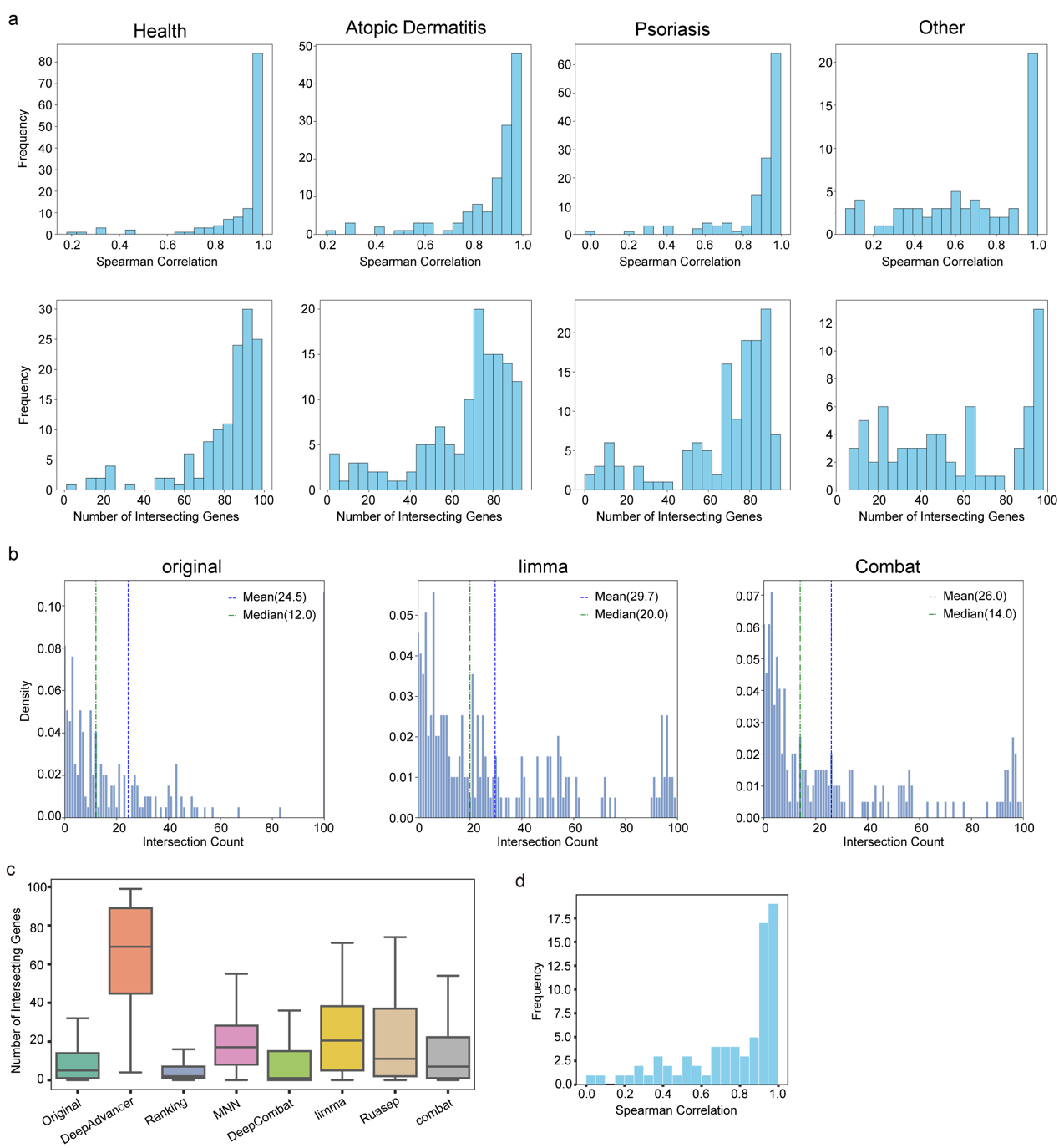

**Supplementary Figure S1:** **Evaluation of expression ratio consistency and gene overlap. a**, Distribution of spearman correlation coefficients and gene overlap counts between DeepAdvancer-corrected and reference expression ratios for each class. **b**, Histogram showing the number of intersecting genes among the top 100 ranked by expression fold-change between each corrected dataset and the reference. **c**, Comparison of intersecting gene counts across different batch correction methods. **d**, Distribution of spearman correlation coefficients between corrected and reference ratios for DeepAdvancer.

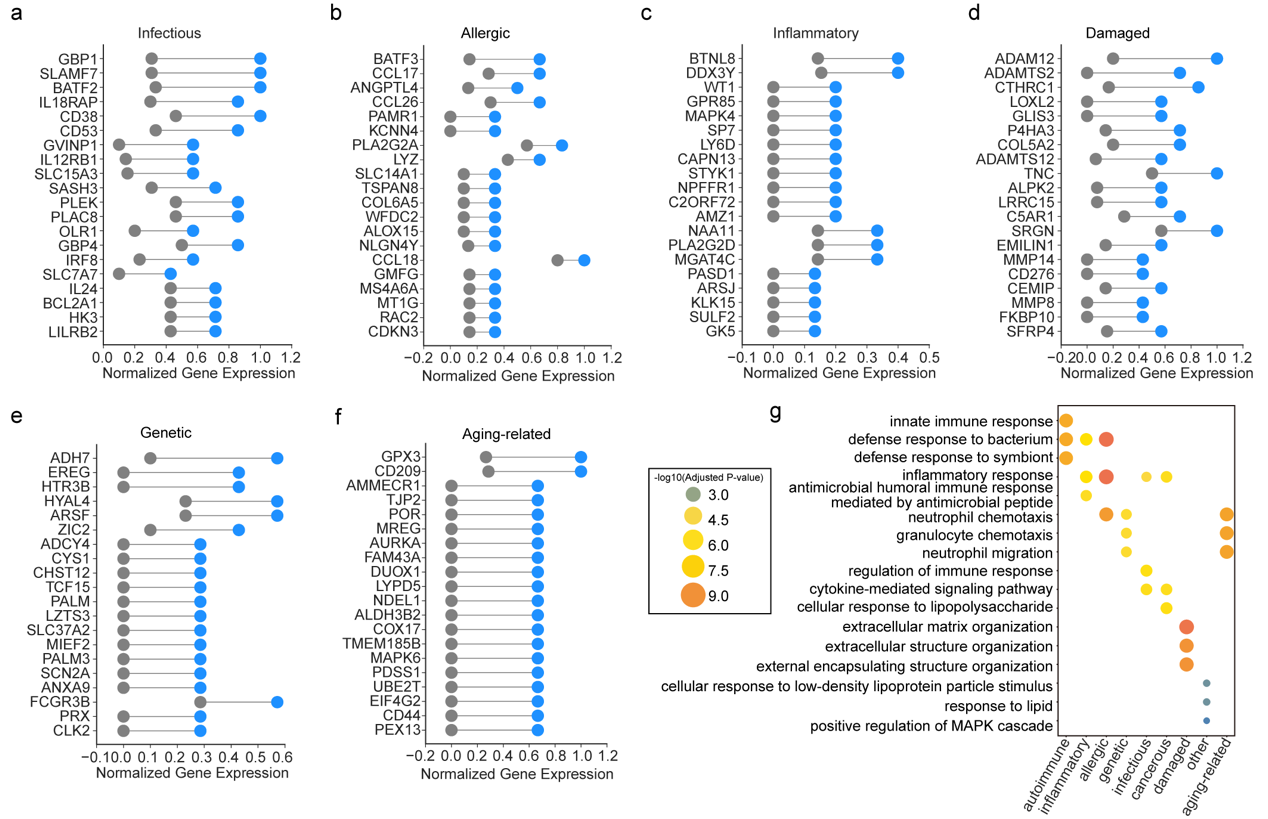

**Supplementary Figure S2: a-f,** The class-specific genes for each class**. g,** The marker pathways for each class.

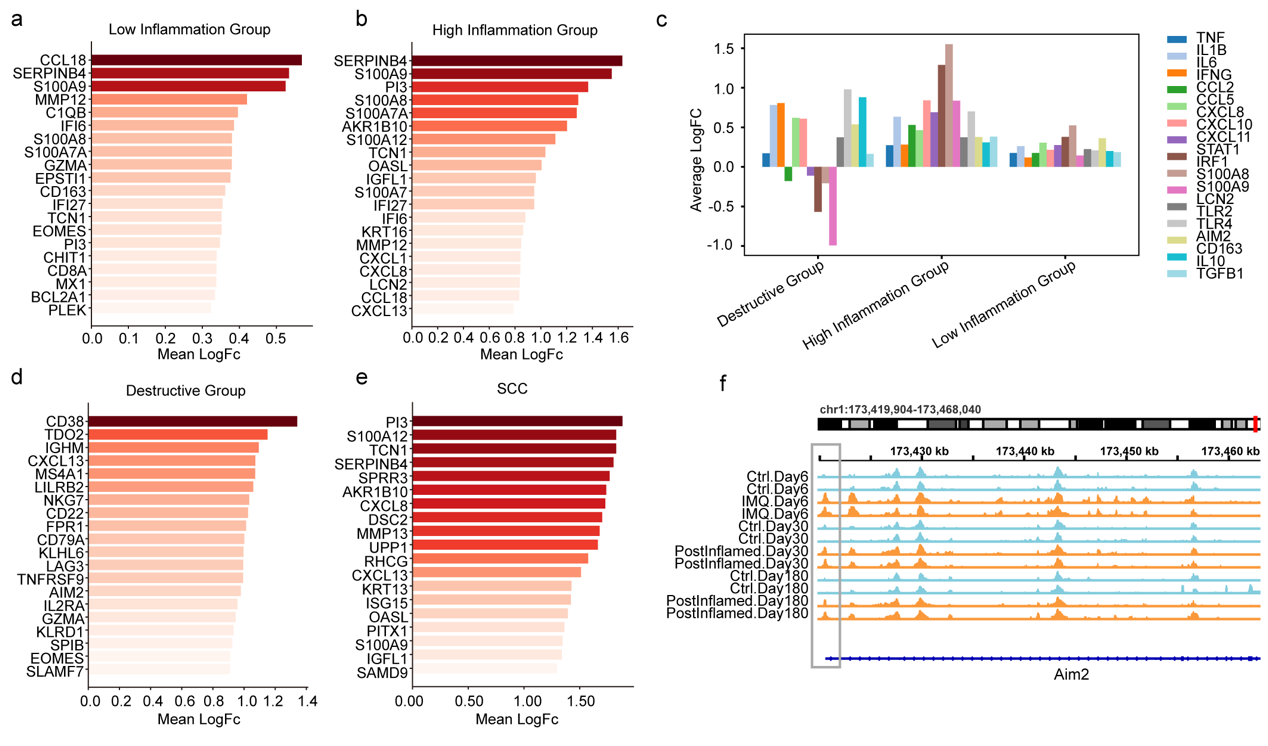

**Supplementary Figure S3: Transcriptomic across skin inflammation states. a–b, d-e, Top genes with elevated expression in the low-inflammation group (a),** **high-inflammation group (b), tissue-destructive group (d), and squamous cell carcinoma (e). c,** Inflammatory **genes are upregulated in high-inflammation group. f, Snapshot of genomic loci whose chromatin-accessible peaks are opened by inflammation at D6 and persist up to 180D following resolution.**

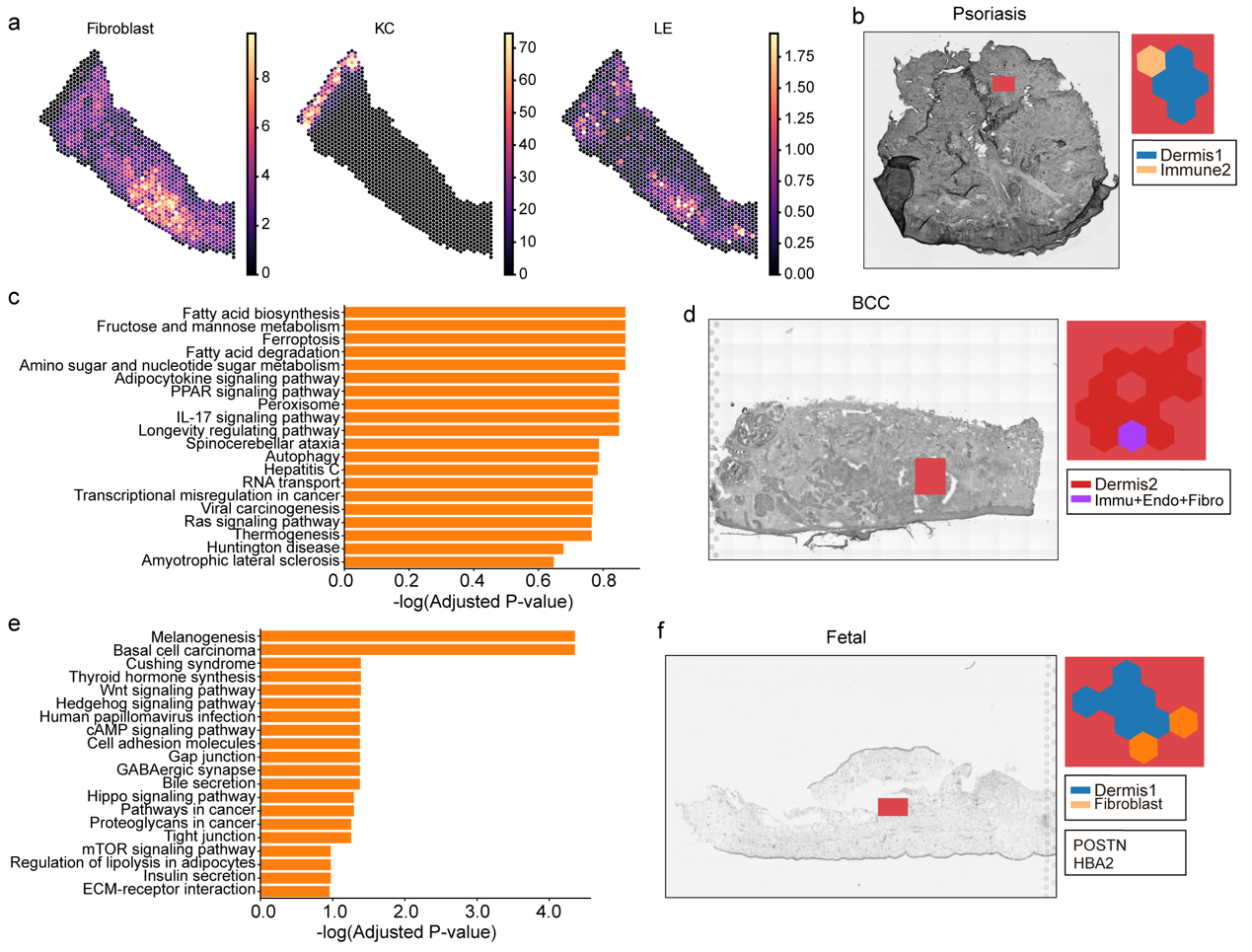

**Supplementary Figure S4:** **Identification and functional profiling of spatial niches in various states. a**, Cell-type composition plot after deconvolution. **b**,**d**,**f,** Spatial niches identified by stNiche in psoriasis (**b**), basal cell carcinoma (BCC, **d**), and fetal skin tissue (**f**). **c**, **e**, Enriched functional pathways associated with the spatial niches in psoriasis (**c**) and BCC (**e**).

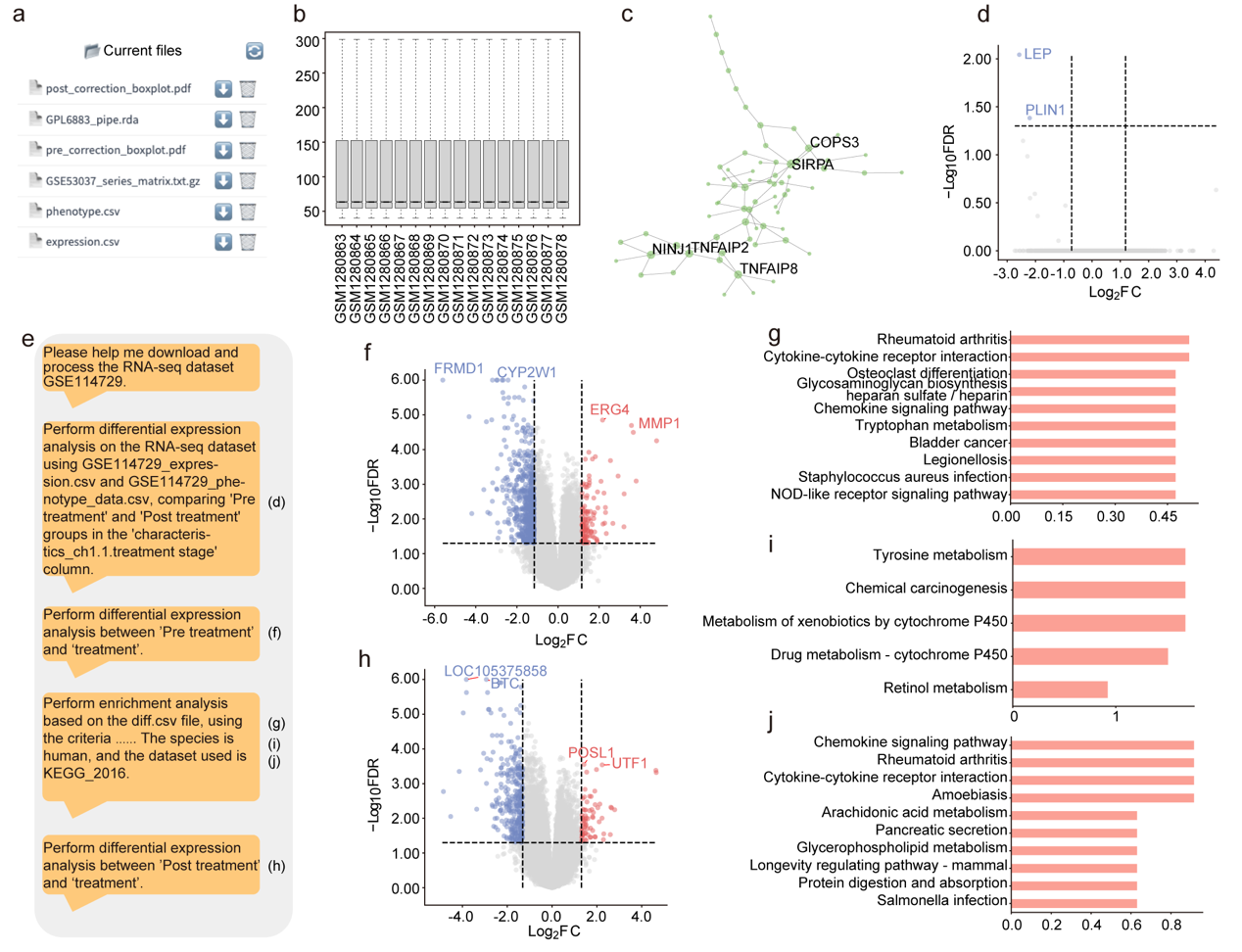

**Supplementary Figure S5: BioinAI-Web supports automated analysis of bulk RNA-seq data. a, Illustration of BioinAI-Web showing result download functions. b, Preprocessed microarray dataset visualized after normalization. c, Module 3 derived from WGCNA analysis on differentially expressed genes from the microarray data. d,** Sequence of **user** prompts **to analyze a bulk RNA-seq dataset and the corresponding results generated by the online platform. e–f, h, Volcano plots showing differentially expressed genes in comparisons between pre-treatment and relapse (e), pre-treatment and post-treatment (f), and post-treatment and relapse (h) samples. g, i, j, Bar plots of enriched functional terms for genes upregulated in post-treatment vs. pre-treatment (g), downregulated in post-treatment vs. pre-treatment (i), and upregulated in relapse vs. post-treatment (j).**

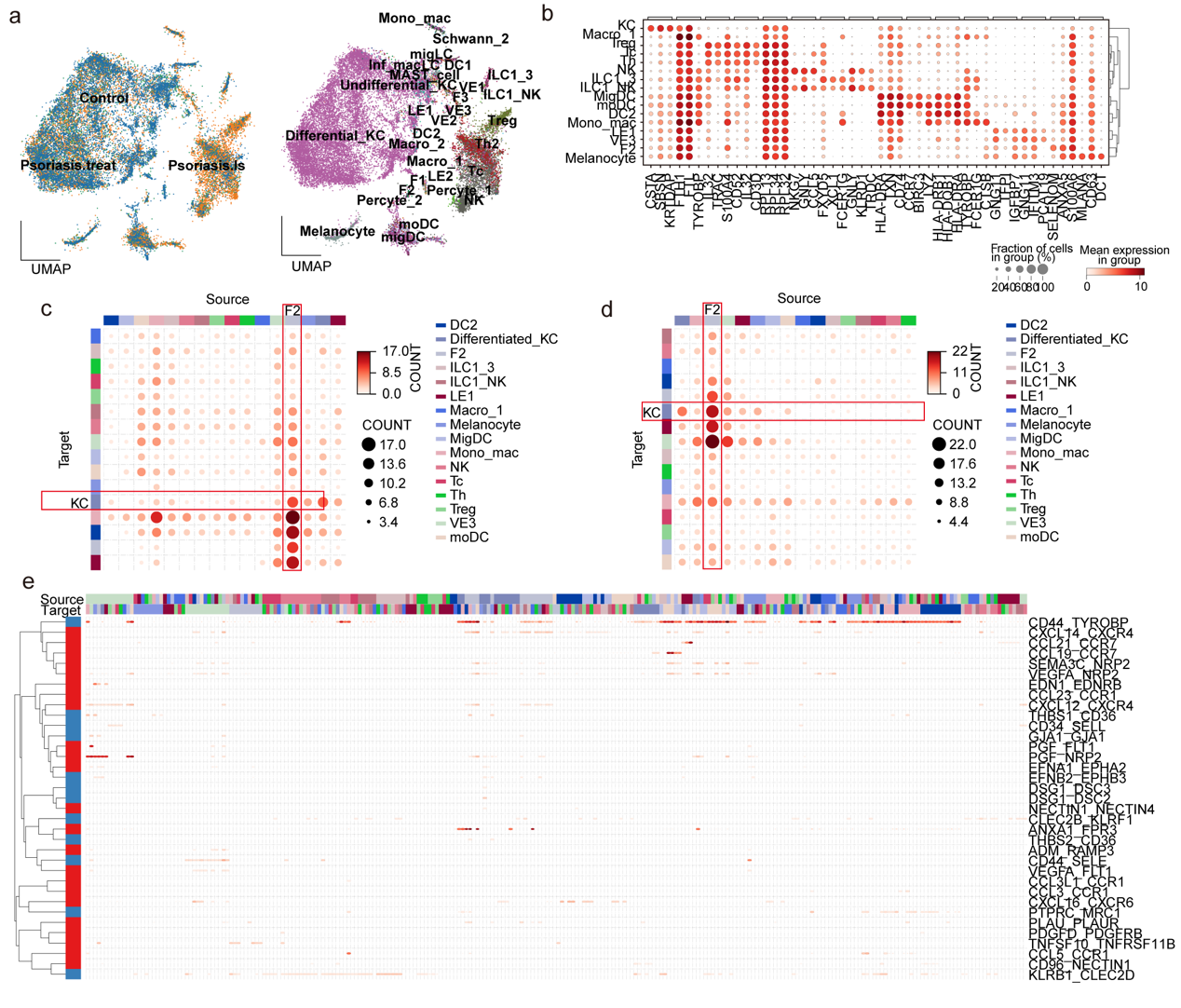

**Supplementary Figure S6: BioinAI-Web enables analysis of single-cell transcriptomic data. a, UMAP plots of integrated single-cell RNA-seq data colored by group (left) and refined cell-type annotations (right). b, Dot plot showing expression of marker genes across subpopulations. c,d, Heatmaps illustrating intercellular communication among subpopulations in healthy controls (c) and post-treatment samples (d). e, Summary of signaling pathways.**
